# Compensation to visual impairments and behavioral plasticity in navigating ants

**DOI:** 10.1101/2023.02.20.529227

**Authors:** Sebastian Schwarz, Leo Clement, Lars Haalck, Benjamin Risse, Antoine Wystrach

## Abstract

Desert ants are known to rely heavily on vision while venturing for food and returning to the nest. During these foraging trips, ants memorize and recognize their visual surroundings, which enables them to recapitulate individually learnt routes in a fast and effective manner. The compound eyes are crucial for such visual navigation; however, it remains unclear how information from both eyes are integrated and how ants cope with visual impairment. Here we manipulated the ants’ visual system by covering one of the two compound eyes and analyzed their ability to recognize familiar views in various situations. Monocular ants showed an immediate disruption of their ability to recapitulate their familiar route. However, they were able to compensate for the visual impairment in a few hours by restarting a route-learning ontogeny, as naïve ants do. This re-learning process with one eye forms novel memories, without erasing the previous memories acquired with two eyes. Additionally, ants having learnt a route with one eye only are unable to recognize it with two eyes, even though more information is available. Together, this shows that visual memories are encoded and recalled in an egocentric and fundamentally binocular way, where the visual input as a whole must be matched to enable recognition. We show how this kind of visual processing fits with their neural circuitry.

**Significance Statement:** If humans look at the world with both eyes, they have no problem to then recognize it with one eye only, and vice-versa. Thus, our way of encoding the world is robust to changes of the visual field. Yet ants do so very differently. Views learnt with two eyes can only be recognized with two eyes, and views learnt with one eye can only be recognized with one eyes (the same eye). However, this rigidity is compensated by a remarkable behavioral flexibility. Upon covering one eye, ants – which can no longer recognize their familiar surroundings – will restart a learning process to store these novel visual inputs in a parallel memory and resume their normal foraging activity.

## Introduction

Self-organized living beings and engineered machines are both able to fulfill their tasks reliably. However, living organisms show flexibility in the way they achieve their functions and can often compensate for unexpected impairments, whereas machines cannot (1). The aptitude for compensation has been well-studied in human with regards to impairments such as cognitive pathologies, aging or brain damage (2, 3), and is evident after morphological impairments, such as when one manages to achieve with one hand what one used to do with two. These forms of compensations require time and likely involve neural rewiring, so-called structural or network plasticity (4, 5)

In insects, which are often assumed to be less versatile than vertebrates (6, 7), the ability to compensate for impairments is usually studied as an instantaneous response and viewed as the product of the evolved natural redundancy and robustness of these systems, rather than the result of neural plasticity through a life time. For instance, the instantaneous change in gait following a single our double leg amputation may be interpreted as a robust and ‘spontaneous’ response of the neural machinery governing leg coordination (e.g., in ants (8), cockroaches (9), stick-insects (10) and other arthropods like crabs (11)). Robustness through redundancy in insects is also well-appreciated in the context of navigation. For instance, representation of directions is based on the integration of a vast array of sensory cues such as visual terrestrial cues (12-15), multiple celestial cues (16-20), olfactory cues (21, 22), wind cues (23, 24), magnetic cues (25, 26) and also self-motion cues (27). Hence, depriving a navigating insect from one modality – or all modalities but one – does not necessarily disrupt their ability to orient. Unilateral suppression of one eye input, however, has a direct impact on the navigational performance of ants (28, 29). Monocular ants may still show evidence of recognition of learnt terrestrial cues but their navigational behavior is drastically affected, revealing an intriguing mix between flexibility and rigidity (28, 29). For instance, monocular desert ants that have learnt a landmark array with one eye only are incapable of recognizing it if the eye cap has been swapped to the other eye, suggesting that there is no inter-ocular transfer of visual terrestrial cues (28). Here again, these studies focused on the insects’ response that immediately follows the manipulation. But whether some plastic, compensatory mechanisms are at play, and given time, can enable a recovery of a functional behavior remains unknown.

Here, we investigated this question by conducting various eye-capping manipulations on visually navigating desert ants (*Cataglyphis velox*). Both, their immediate response as well as the potential compensatory effects emerging after a longer period of time were observed and analyzed. The results show that visual manipulations via eye-capping caused substantial impairments that disrupt the ant’s ability to home. However, a few hours upon the visual impairment, ants recovered a functional navigation behavior, indicating the existence of a profound plasticity and behavioral flexibility. We explored the mechanisms underlying this plasticity and, as a corollary, gained insight into the way visual information is stored in their brain.

## Results and Discussion

### One-eye-capping disrupt learnt route-following

Iberian desert ants (*Cataglyphis velox*) were individually marked and let free to navigate back and forth along an 8.0 m long route between their nest and a feeder (Fig. 1A). The surrounding natural landscape provided plenty of visual information and the route floor was covered with a 1.2 m wide wooden board ensuring an even substrate for the navigating ants. Once experienced to the route (after at least 10 foraging trips), ants were captured at the feeder one by one for the eye-cap treatment: either their left or right compound eye was covered with opaque paint. The eye-capped ant was then provided with a food item, released near the feeder and her homing path was recorded. Regardless of the side of the eye cap, ants failed to navigate directly toward their nest. Instead, their initial direction showed a bias toward the side opposite to the eye cover (Fig. 1B) as previously reported in Tunisian desert ants (28). All eye-capped ants (10 out of 10) veered sidewise and fell off their usual route corridor, defined by the wood plank on the floor (Fig. 1A). To enable them to reach home, ants falling off the wood boards were systematically captured and re-released back onto the route beeline (Fig. 1A, dashed line) where path recording resumed. Eye-capped ants veered off the boards 3.42 times on average before reaching their nest surrounding, where recording was stopped (Fig. 1A, Fig. SI1). The difficulty to head home was obvious along the complete homing path (Fig. 1A). By comparison, Sham ants, which received a paint mark on the head but untouched compound eyes and ocelli, showed clearly oriented (Fig. 1B) and much straighter paths (Fig. 1A, Fig. SI1) and almost never left their route corridor (1 out of 17 ants fell one time, so a probability of 0.06 time/ants on average). Thus, covering one compound eye drastically affects navigation in homing ants.

**Fig. 1.**
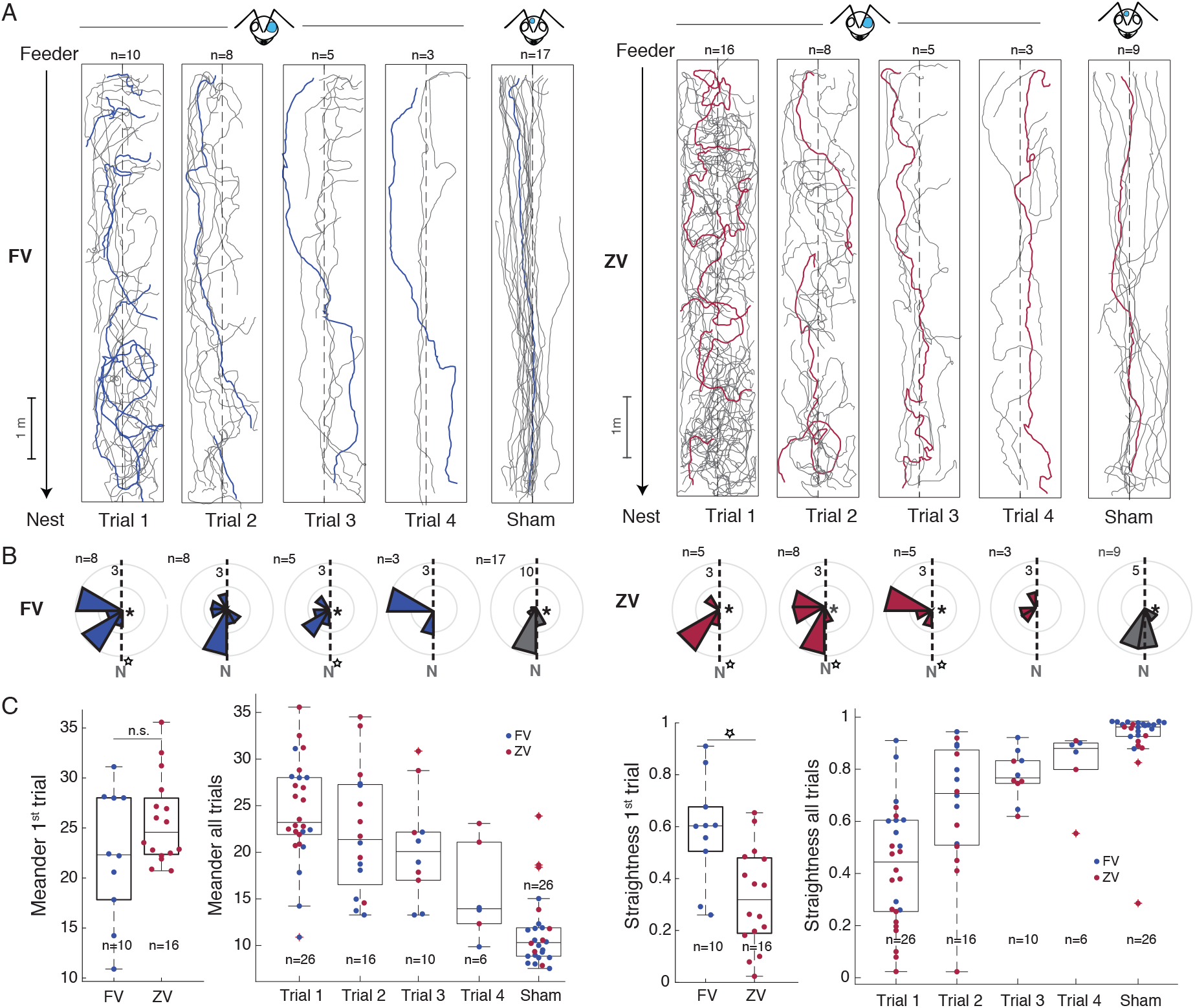
Initial side biases of homing paths in eye capped ants. Black arrows: travel direction of eye capped ants from feeder to nest along a straight foraging route; dashed line: middle of the foraging route; grey lines: ant paths, gaps between paths occurred when ants ran off the board and were re-released in the middle of the homing route; ant head sketch: eye cap condition (this applies to all other Figures). Left- (LEC) and right (REC) eye capped ants were pooled. Paths of REC ants were mirrored and depicted as LEC ants (painted blue eye in ant head). (A) Homing paths from trial 1 to 4 in full vector- (FV) and zero vector ants (ZV) together with homing paths of Sham ants with uncovered eyes. Side biases in paths are visible in both FV (blue example paths) and ZV (red example paths) ants but not in Sham controls. (B) Dashed line: feeder nest direction; number at the outer rim of diagrams: number of ants per sector. Circular histograms depicting directional distribution of ant’s paths after 0.2 m of travel across trials and condition. Significant differences from random distributions are indicated with a star (Rayleigh test) at the center of the diagrams and significant differences from the nest direction (N) are indicated with and open star (S-test, for statistical details see Table S1). (C) Meander and straightness of ant paths across trails and condition. FV and ZV vector ants differ significantly in the level of straightness in their first homing path but not in the level of meander. With increasing trial number both FV and ZV paths resemble more Sham paths with low meander and high level of straightness.

The compound eyes of ants extract information from both, celestial compass cues — which are key for path integration (PI) — and terrestrial cues — which are key for learnt views during route-following (30). To test whether the behavioral defect observed in eye-capped ant is due to a disruption of the PI system or the use of learnt views, the experiment was repeated by using this time so-called zero-vector (ZV) ants. While full-vector (FV) ants are captured at the feeder, ZV ants were captured on their way home just before entering their nest, then received an eye-cap and were released with their food item right near the feeder. The PI vector of a ZV ant does no longer point toward the nest; hence the ant can solely rely on learnt terrestrial cues for homing (31).

As for FV ants, covering one eye of ZV ants strongly disrupted their ability to navigate home (Fig.1A). Their initial directions were also biased toward the open eye side (Fig. 1B) and all of them (16 out of 16) repeatedly ran off their usual route corridor (4.43 times/ant on average) to the contrary of ZV sham ants (0.33 times/ant on average). Their overall paths were significantly less straight (Anova: *F*_*1,24*_ = 10.7, *P* = 0.003) but only marginally more meandrous (Anova: *F*_*1,24*_ = 2.42, *P* = 0.133; Fig.1C) than eye-capped FV ants. Consequently, eye capping strongly impairs the use of learnt terrestrial cues and the directional input provided by the PI is helping only slightly the FV eye-capped ants to maintain straighter paths. This is in line with previous evidence that information based on celestial but not terrestrial compass cues undergoes inter-ocular transfer (28).

### Eye-capped ants spontaneously recover their route-following behavior

Surprisingly, within a few hours, some of the tested eye-capped ants reoccurred at the feeder again, which provided the opportunity to record their subsequent homebound trips. With increasing homing trials, eye-capped ants gradually recovered their navigational efficiency (Fig. 1). To ensure this recovery was not simply due to an increased reliance on PI, eye-capped ants reaching their nest were captured as ZV ants and released again at the feeder for a second run home. Whether as FV- or ZV ants, their initial direction upon release still tended to be biased toward the open eye side until the 3^rd^ homing trial (Fig. 1B; see Table S1 for circular statistics). However, both FV- and ZV paths showed progressively less local meander (Anova: *F*_*0*.*1,49*.*4*_ = 25.170, *P* < 0.001) and more overall straightness (Anova: *F*_*1*.*1,51*.*3*_ = 32.52, *P* < 0.001; Fig. 1C). During the 4_th_ homing trial the path straightness of eye-capped ants resembled the one from the sham ants (Anova: *F*_*1,30*_ = 2.68, *P* = 0.112), and even though they still showed slightly more local meandering (Anova: *F*_*1,30*_ = 5.39, *P* = 0.027; Fig. 1C), most eye-capped foragers (5 out of 6) managed to home while no-longer exiting their route corridor. Within a relatively short time period, eye-capped ants can thus recover the ability to follow their familiar route again, using terrestrial cues. These insects are therefore able to compensate impressively fast the strong impairment caused by losing the visual input of one eye, showing a remarkable plasticity in their navigational capacities.

### Eye-capped ants show a persistent lateralized sensory-motor defect

The fact that the initial direction of both FV- and ZV ants upon eye-covering was biased toward the open eye side as compared to the correct route direction (Fig. 1B) provides two insights. First, because ZV ants did not head randomly (or backtracked) like they do on visually unfamiliar terrain ((32), Fig. 2B), freshly eye-capped ants must still be able to derive some information from their visual route memory. Second, because the side of the directional bias depends purely on the side of the capped eye (Fig. 2A, C), a lateralized sensory-motor defect caused by losing one-eye input might be at play.

**Fig. 2.**
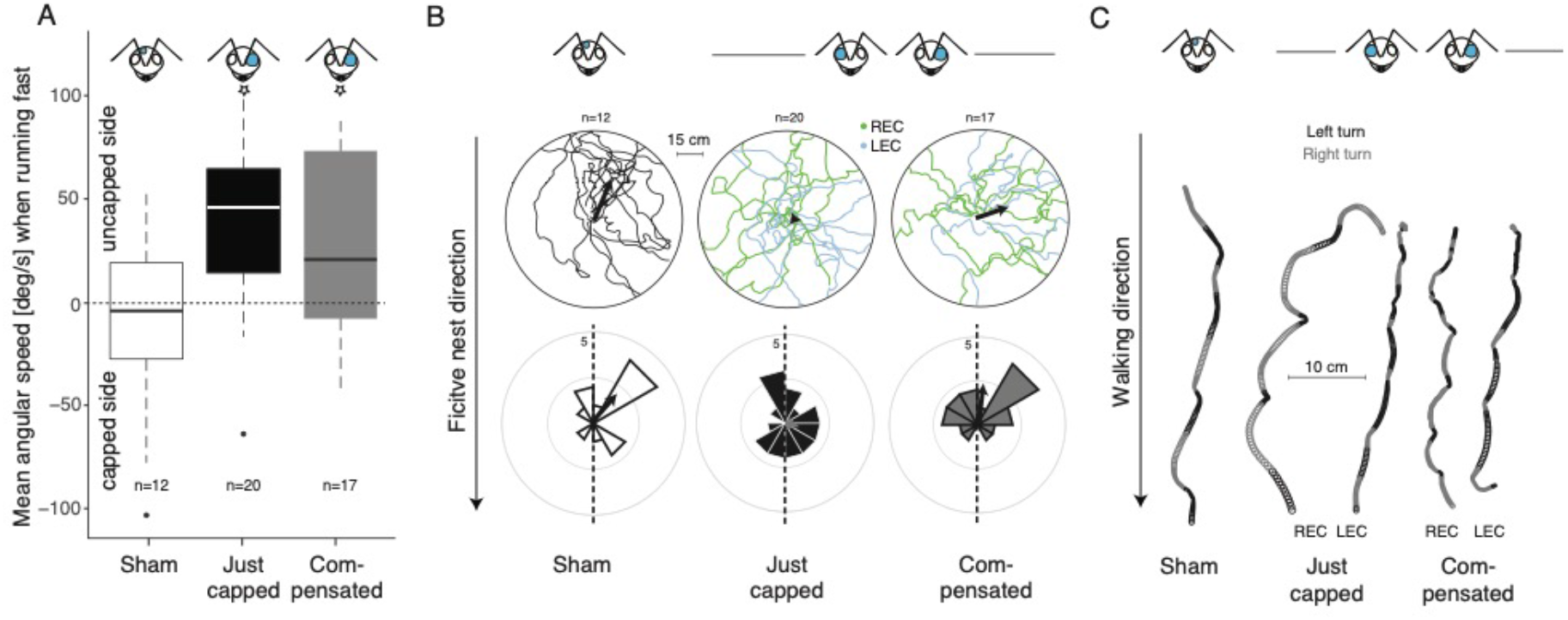
Initial side bias in eye-capped ants released in unfamiliar surroundings. (A) Distribution of side preferences of ants with just capped eyes (Just capped) and ants that had compensated (Compensated) for the visual impairment in their familiar foraging route. The characteristic motor deficit, displayed by capped ants toward the side bias of the uncapped eye is evident in Just capped ants, as represented by the higher mean angular speed when turning to the uncapped eye side. Compensated ants show a lower side bias and resemble more the even distribution of Sham ants. Stars indicates significant difference to random, one-sided Wilcoxon test (see text for details). (B) Tracked paths of Sham, Just capped and Compensated ants (upper panels) and initial (60 cm radius) heading directions (lower panels). Black arrows depict mean heading directions. (C) Example paths of ants with (Just capped, Compensated) and without (Sham) the characteristic motor deficit caused by eye-capping.

To investigate this lateralized sensory-motor defect independently of the expression of the recognition of visual memories, freshly eye-capped ants were released in unfamiliar surroundings and their behavior was analyzed based on video recordings. Eye-capped ants tended to regularly alternate between brief bursts of speed when turning toward the open eye side and pauses by rotating on the spot toward the covered eye side (Fig. 2C). This can be simply quantified by assessing whether turning direction is biased on one side when the ants forward speed is above the individual’s average. While sham ants with both eyes open showed no bias (Sham ants: one-sided Wilcoxon test: V = 32, *P* = 0.715, Fig. 2A) freshly one-eye capped ant turned more toward the open eye when walking fast (Just-capped ants: one-sided Wilcoxon test: V = 188, *P* = 0.001; Fig. 2A).

To test whether the observed recovery of route-following behavior is due to a compensation over time of such a lateralized sensory-motor defect, we tested whether the bias persisted in ants after they had recovered their route-following behavior with one eye. Eye-capped ants that had recovered their route still displayed the lateralized defect on unfamiliar terrain (Compensated ants: one-sided Wilcoxon test: V = 113, *P* = 0.044, Fig. 2). We noted that, given longer period of time with the eye cap, this bias eventually tended to fade (data in preparation) showing the ability of experienced ants to also correct for such a sensory-motor defect. However, the persistence of the lateralized bias here (Fig. 1, 2) shows that the ant’s route recovery is due to a process that is quicker and different from overcoming such a sensori-motor defect. Also, this longer lasting sensori-motor defect may explain why one-eye ants having just recovered their route still showed more meandering than sham ants (Figure 1).

### Eye-capped ants compensate by re-engaging in a new route ontogeny

We next investigated whether the recovery of route-following behavior in eye-capped ants is based on the ability to eventually recall previous binocular memories, or alternatively, based on the formation of novel, monocular route memories. To do so, we covered the left or right eye of a new cohort of experienced, individually marked ants, and released them back to their nest. Their behavior was recorded once they emerged outdoors again. Upon leaving their nest entrance these freshly eye-capped ants behaved remarkably similar to naïve foragers exiting their nest for the first time: they displayed typical learning walks (Fig. 3). These convoluted paths and pirouettes enable them to expose their gaze in multiple directions for visual learning (33). Learning walks are likely a consequence of perceiving an unfamiliar scenery when leaving the nest. Indeed, experienced ants may also display a few zigzags and pirouettes upon departure if an alteration of the visual surrounding has occurred around the nest; but in general, they rapidly scoot along their familiar outbound route again (33, 34). Here, the eye-capped ants remained at first very close to the nest and re-entered their colony often, like naïve ants do (Fig. 3C). They displayed on average more than 15 subsequent learning trips before reaching the familiar feeder located m away (Fig. 3A, B) supporting the idea that the scenery appeared strongly unfamiliar to these eye-capped ants. Interestingly and contrary to naïve ants, learning walks of freshly eye-capped ants were biased toward the feeder direction (Fig. 3A); this was true from the first learning walks onward (Fig. 3C, One-tailed T-test: *P* = 0.019) indicating that previous memory of the feeder direction persisted despite the eye-cover. Whether this directional memory was due to remnant memories of terrestrial cues learnt with both eyes, or the expression of a stored food-ward celestial compass vector (35, 36) could not be disentangle here.

**Fig. 3.**
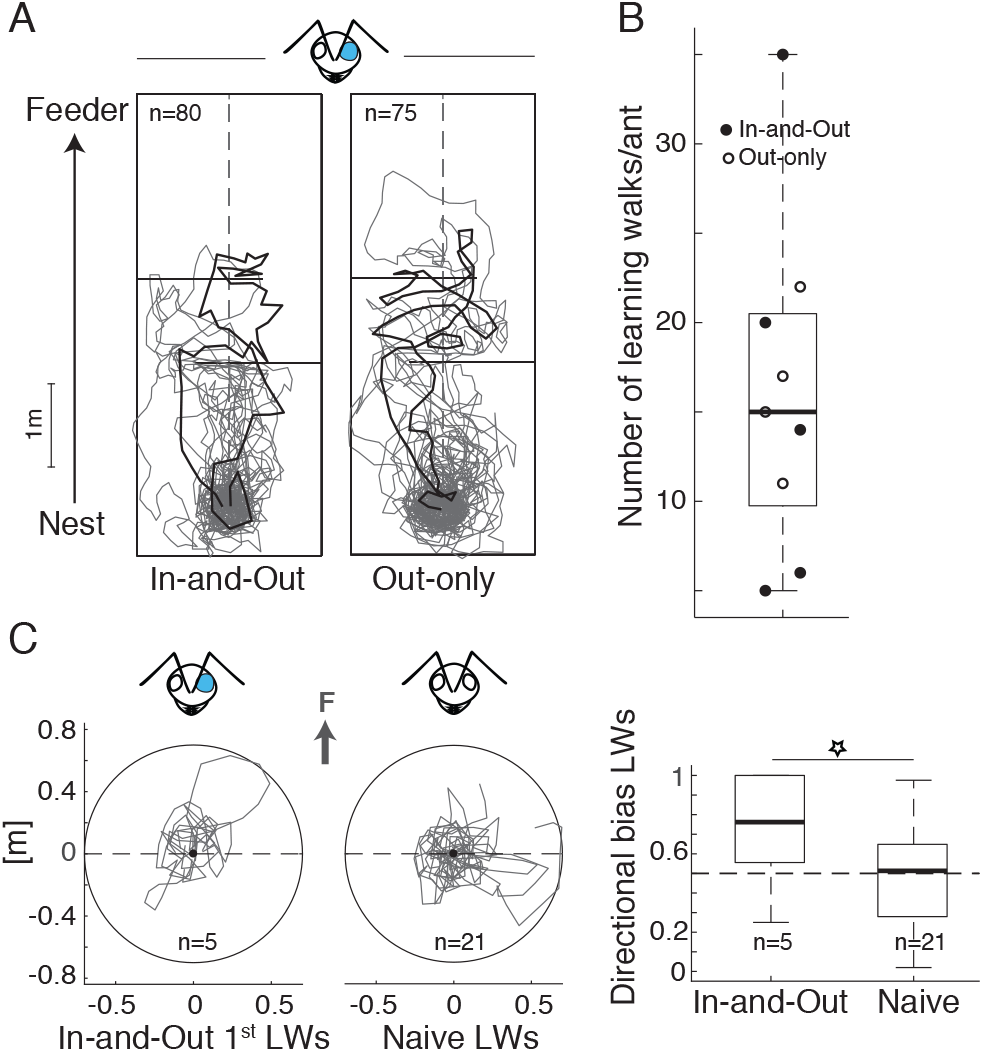
Learning walks (LWs) in eye capped ants. Horizontal lines: one-way baffles on the foraging route that could be traversed during outbound trips but had to be negotiated during inbound trips (this also applies for Figs. 4 and 5). (A) LWs displayed by In-and-Out and Out-only ants during training. Black bold lines: example paths. (B) Distribution of the number of LWs in In-and-Out and Out-only ants. (C) Comparison between directional path bias of first LWs in In-and-Out and naïve ants. In-and-Out ants show a significantly higher bias toward the feeder direction (grey arrow) than naïve ants.

The subsequent out- and inbound (i.e., homing) trips of these eye-cap ants were also recorded, which, as expected, showed gradual improvements (Fig. SI3 and SI4). After 8 successful trips between the nest and feeder, all the recorded eye-capped ants (In-and-Out) had fully recovered their ability to run between the nest and the feeder without colliding into baffles (Fig. 4A). Tested as ZV ants, these individuals could home equally well, (Fig. 4A) showing as previously, that they used terrestrial cues.

**Fig. 4.**
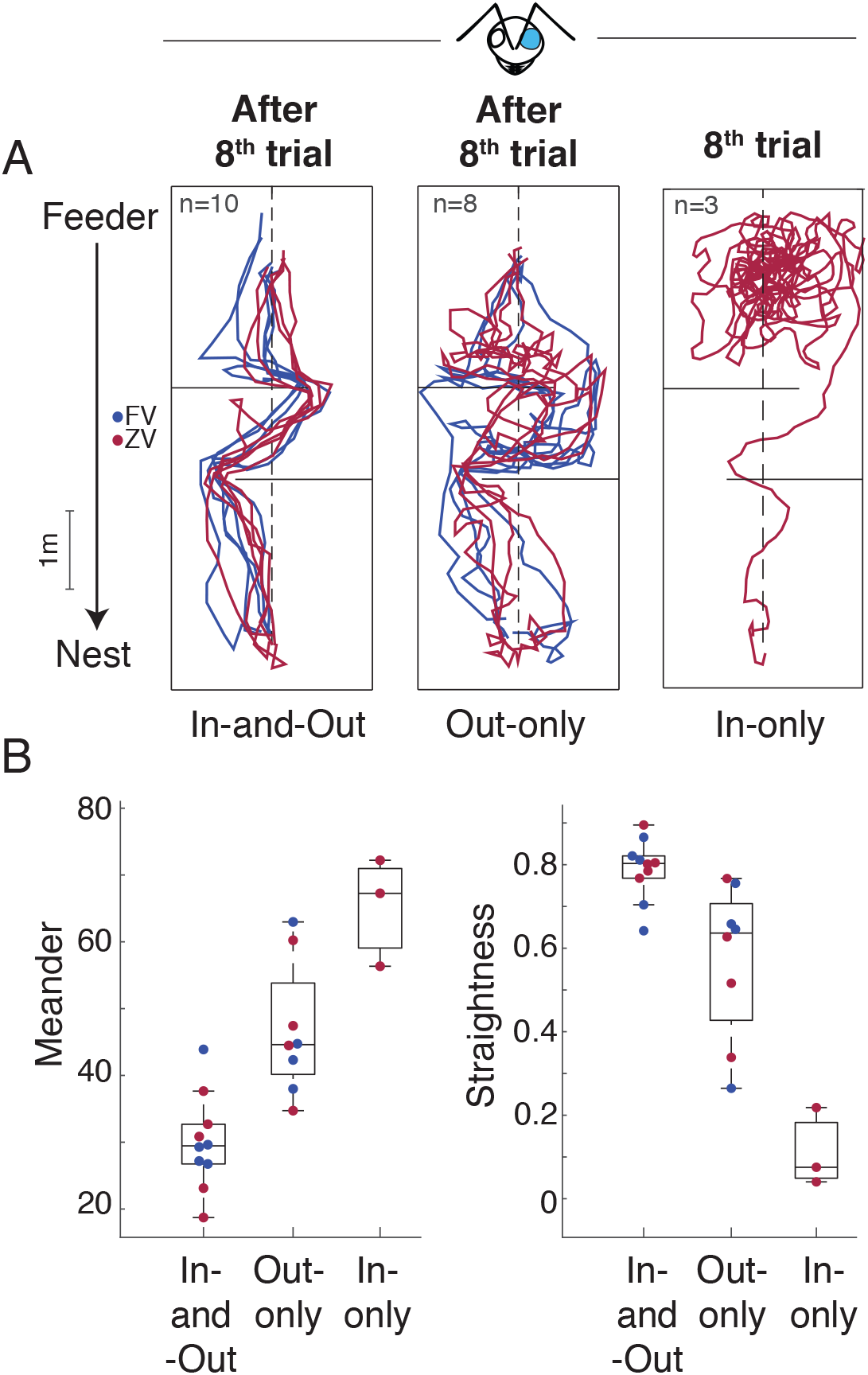
Paths and path characteristics of eye capped ants after compensation with in and outbound trip (In-and-Out), no inbound trip (Out-only) and no outbound trip (In-only). (A) Paths of homing ants after 8 training trials. All In-and-Out and Out-only ants were able to home whereas In-only ants struggled and only one individual succeeded. (B) Meander and straightness of homing paths after 8 training trials. In-and-Out ants displayed paths with the lowest meander (Anova: In-and-Out vs. Out-only: *F*_0.29,0.07_ = 4.122, *P* = 0.001; In-and-Out vs. In-only: *F*_0.63,0.11_ = 5.981, *P* < 0.001) and highest level of straightness (Anova: In-and-Out vs. Out-only: *F*_-0.02,0.06_ = -3.436, *P* = 0.003, In-and-Out vs. In-only: *F*_-0.69,0.09_ = -7.327, *P* < 0.001), followed by Out-only and In-Only ants.

To ensure that this recovery was actually due to performing learning walks, and not simply due to the time passed while navigating outdoors, the experiment was replicated by using two additional cohorts of freshly eye-capped ants that had previous experience (with both eyes) of the route. In one group (In-only), eye-capped ants were systematically captured upon exiting the nest and released at the feeder for homing, preventing them to display learning walks around the nest and outbound trips to the feeder. In the other group (Out-only), eye-capped ants were free to display learning walks but upon reaching the feeder, these ants were systematically captured and released inside their nest entrance, preventing them to perform their homing runs (inbound trips). This latter group of ants (Out-only) displayed similar learning walks than the previous condition (Fig. 3), and after 8 successful outbound trials up to the feeder, foragers were able to home quite well, albeit not as successful as ants that had experienced both out- and inbound trips (Fig.4 and Fig. SI3). This shows that inbound trips are helpful but not crucial for route recovery. Contrastingly, ants deprived of learning walks (In-only) showed no improvement in their homing ability despite spending a long time navigating outdoor (Fig. 4). On the contrary, they showed a decrease in homing performance across trials (Fig. SI3), suggesting again that freshly eye-capped ants have strongly impaired but remnant binocular visual memories of the route, but that subsequent experience outdoor does not enable to re-access them, or form new functional ones. Note that In-only ants, as being transferred from the nest to the feeder, could not rely upon path integration. After eight attempts of homing without learning walks, these ants searched at length around the release point (feeder) and only one individual eventually managed to home (Fig. 4). It was rather difficult to conduct this condition, as In-only ants mostly failed to reach their nest during training. After 10 min of search and an unsuccessful inbound trip the foragers had to be put back manually to the nest. Unfortunately, many of those individuals stopped their foraging activity and hence could not be tested anymore.

In sum, performing learning walks and outbound trips is crucial for eye-capped ants to recover their route. This echoes what is observed in naïve ants with untouched eyes, for whom learning walks and outbound trips are key for learning an inbound route, while inbound experience may help but seems to be insufficient on its own (37-41). Together, this suggests that eye-capped ants cannot recognize the learnt route with both eyes to full capacity and thus perceive at first the world as quite unfamiliar. This unfamiliarity triggers learning walks upon leaving their nest, which enables them to form novel memories of the terrestrial cues and eventually re-learn the route monocularly. Therefore, ants with previous experience possess the flexibility to recapitulate the natural ontogeny of route learning for a second time.

### Ants learn binocular visual memories

Previous behavioral work in ants and bees has shown that visual memories learnt with one eye cannot be retrieved using the other eye, suggesting that these insects form two separated visual memories for each eye, with an absence of inter-ocular transfer between these visual memories (28, 42). The current study revealed that visual memories acquired with both eyes cannot be retrieved with one eye. Rather than two separated visual memories for each eye, one possible explanation is that visual memories are fundamentally binocular, that is, their recall implies the correct and simultaneous combination of both left and right visual input. Indeed, neurobiological studies show that each eye sends bilateral visual projections to the Kenyon cells (KCs) in both the left and right hemispheres of the mushroom bodies (MBs)(43), where visual memories for route-following are formed (44-46). What’s more, visual projection to the KCs are pseudo-random, therefore, it seems quite likely that individual KCs, whether in the left or right hemisphere, receives input from both eyes, and thus must receive the correct bilateral input to be activated (Fig. 6). If this hypothesis is correct, it predicts, rather counterintuitively, that memories acquired with one eye could not be retrieved with both eyes.

To test this prediction, a novel cohort of ants from a new nest was eye-capped either on the left or the right eye. Crucially, the training to the foraging route and the transformation of the visual scenery by clearing the floor and altering the natural bushes and other terrestrial cues around the route was done afterward. The set-up was similar to the former experiment: a 5.0 m long route containing two baffles in the middle and one feeder at the end. These ants had binocular memories of the previous natural surroundings but experienced the foraging route and its novel surroundings only with an eye-cap (i.e., monocular). After a few hours, the eye-capped ants were familiarized with the new surroundings, discovered the feeder and learnt to home successfully along the novel route (Fig. 5A). When captured at the nest and released again as ZV ants near the feeder, these now experienced eye-capped foragers had no difficulties to recapitulate the route, confirming that they had formed (monocular) visual memories of the terrestrial cues (Fig. 5A, left panel). However, when uncapped, that is, when exposed for the first time to this particular route with two eyes, the ants struggled immensely to home (Fig. 5A, right panel). All uncapped ants searched predominantly in the upper section of their foraging route incapable of negotiating the baffles, only 2 out of 10 eventually managed to reach the nest entrance (Fig. 5A, right panel). Even when some (5 out of 10) of these ants were re-released beyond the baffles, closer to the nest, they were not able to home successfully and wandered around seemingly without purposeful orientation (paths not shown). Thus, ants with two eyes were unable to recognize the route learnt with one eye. In other words, adding the visual input of the second eye prevents the access to the visual memories acquired with one eye only. Therefore, when it comes to navigation, visual memories are binocular.

**Fig. 5.**
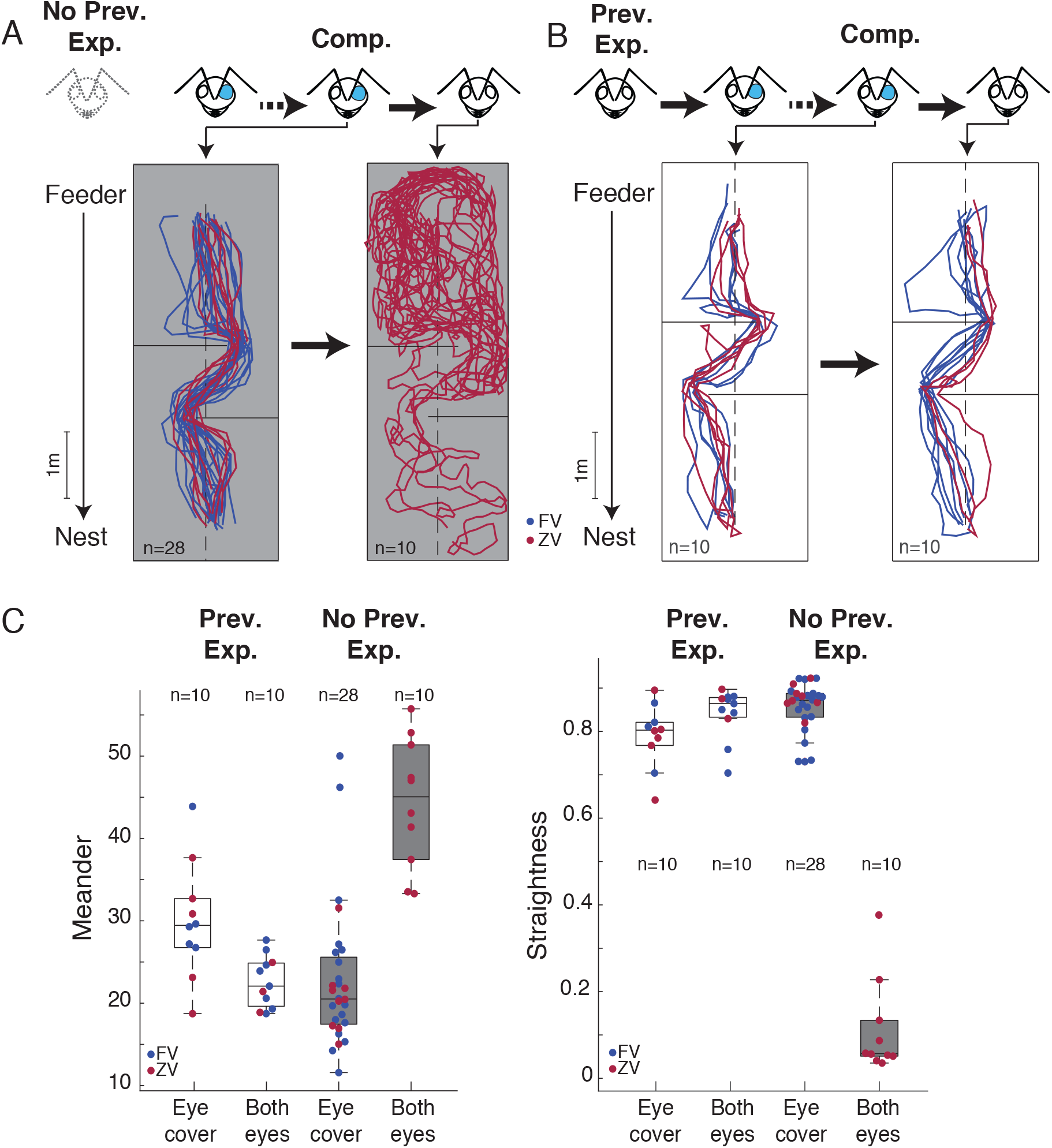
Visual impairment before route learning. (A) Ants that learnt the route with one eye covered, hence no previous experience (no prev. exp.) of the route with uncovered eyes. After compensation (comp.) and reaching the feeder, homing paths are displayed with the eye cap still in place (left panel) and with uncapped eyes (right panel). Ants without previous experience of the route struggle to find back to the nest once the eye cap is removed. (B) Ants that learnt the route with both eyes, hence previous experience (prev. exp.) of the route before eye capping. After eye capping and reaching the feeder, homing paths are displayed with the eye cap still in place (left panel) and the eye cap removed (right panel). Ants with previous experience of the route have no difficulties to find back to the nest. (C) Path characteristics of ants with and without previous experience of the route before eye capping. Ant with previous experience show paths with low meander and high level of straightness after eye cap was removed whereas ants without previous experience of the route show paths with high meander and low level of straightness with both eyes uncovered (Anova meander: *F*_-0.13,0.05_ = -2.487, *P* = 0.018, Anova straightness: *F*_0.07,0.02_ = 3.281, *P* = 0.002).

In terms of neural implementation, this supports the idea that many KCs in the MB receive input from both eyes simultaneously (Fig. 6). It remains likely however that some KCs receive input from one eye only, which could explain why some freshly eye-capped ants, or freshly uncapped ants, even though strongly impaired, could still derive a rough estimation of the nest direction, as if they recognized the scene only partially or sporadically. The variation in homing performance observed across individuals may be explained, at least in part, by the more or less fortunate random connectivity in their mushroom body.

**Fig. 6.**
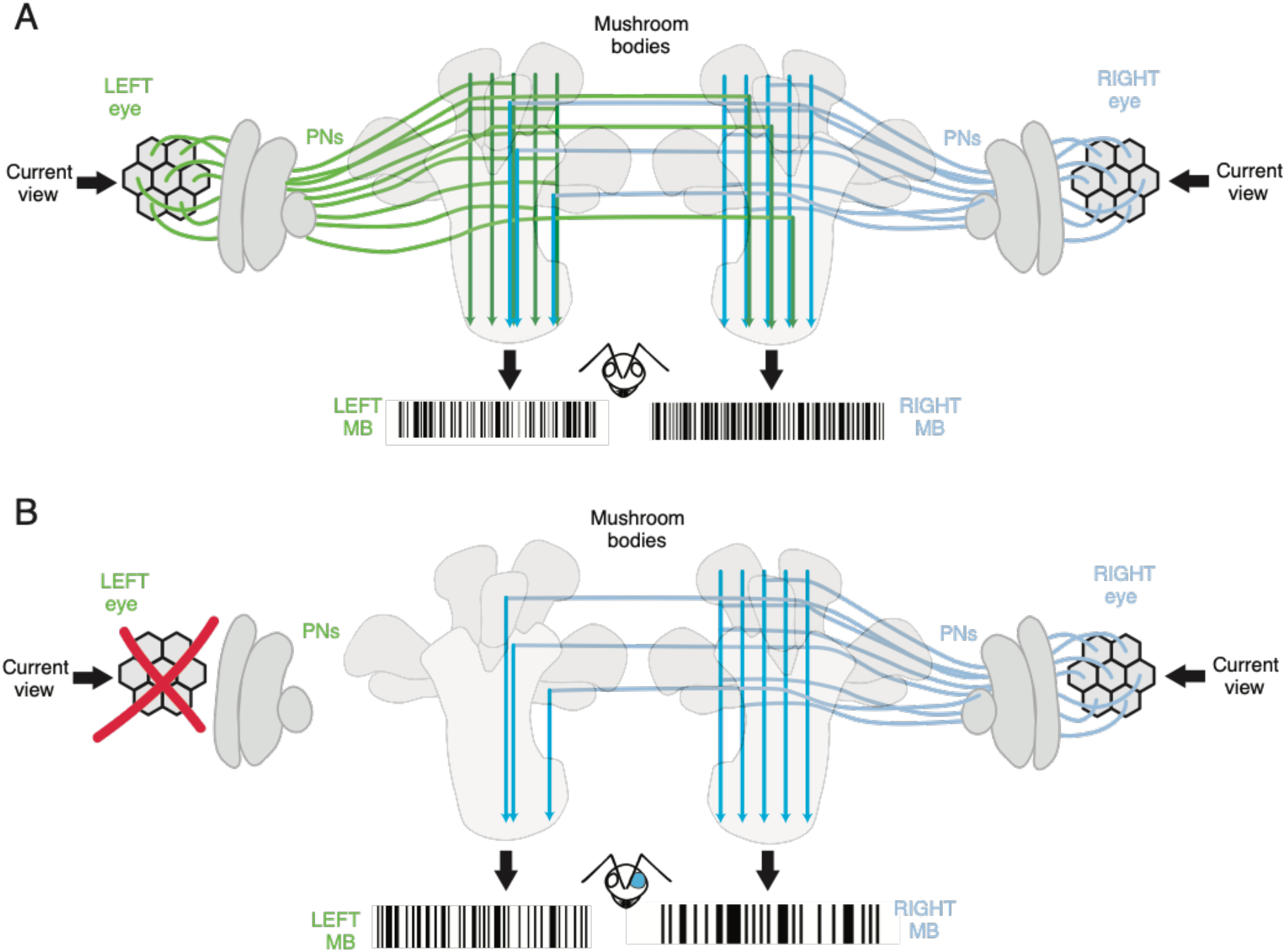
Simplified model depicting neurobiological implications. (A) Both compound eyes feed into both Mushroom Bodies and elicit via projection neurons (PN) a certain set of Kenyon cells for each perceived view along a familiar route. (B) If the left eye is impaired (eye cap) the patterns of Kenyon cells are disrupted as they perceive only input from the uncapped right eye. Hence, the familiarity of the visual scenery is altered and homing ants need to compensate for the lack of visual information by relearning the surrounding scenery. The different bar codes represent different activated sets of Kenyon cells for a given view, for each compound eye.

### Eye-cap ants do not forget past binocular memories

To control whether the visual impairment observed with freshly uncapped ants (Fig. 5A, right panel) was indeed due to an inability to access visual memories sorted with one eye and not just an inherent consequence of the recent recovery of bilateral visual input, the previous experiment was re-run with a cohort of ants that, this time, had previous two-eyed experience of the route. These ants were eye-caped, let free to re-learn the route with one eye, then ‘uncapped’ and tested. Contrary to the previous cohort of ants with no previous experience of the route, these foragers did not struggle whatsoever to home toward the nest with uncapped eyes, even when tested as ZV ant (Fig. 5B). This shows that uncapping the ants bears no inherent issue and thus confirms that memories acquired with one eye are no longer accessible with two eyes (Fig. 5A). In addition, it shows that the latter cohort of ants had not forgotten their former visual memories of the route acquired with two eyes. Learning the route anew with one eye does not override the memories of the previous two-eye memories.

In neurobiological terms, one-eyed memories and two-eyed memories are likely to recruit different set of KCs in both hemispheres, with perhaps a certain amount of overlap between them (Fig. 6). This can be viewed as learning and remembering two different routes, which desert ants can also do (47). Indeed, ants possess hundreds of thousands of KCs (43) and models of the MBs show that this offers memory space for recognizing a large amount of visual sceneries (48, 49); enough to remember views around the nest, along multiple routes or as shown here, along the same route but with monocular and binocular inputs.

## Conclusion

The response of navigating ants to visual impairments shows a surprising mix between rigidity and flexibility. Rigidity in the sense that the recognition of visual memories is not robust to changes in the visual field, highlighting a fundamental difference with the way human encode the visual world. Flexibility in the sense that experienced ants manage to compensate such a deficit by re-engaging a route learning process in a similar way than naïve ants do at the onset of their foraging ontogeny. With time, ants re-learn and recognize a route with ease, and the newly acquired memory does not override previous ones. Investigating the plastic neural mechanisms underlying these feats will form a great agenda for future research.

## Material and Methods

Experiments took place during June and July 2017-19 on a plain open field with grassy vegetation close to the harbor in the metropolitan area of Seville, South of Spain. Three different nests of the Iberian desert ant *Cataglyphis velox* were used for training and testing. Workers exhibit behaviors typical for solitary foraging ants that venture out of the nest to find food without the help of pheromone trails (50). Instead their navigational guidance is primarily based on visual input derived from celestial and terrestrial sensory cues (51).

### General experimental set-up

All set-ups shared a similar basic design, which is described in the following while specific differences were appropriately mentioned above. Ants were trained to follow a route from the nest to a feeder that provided food *ad libitum* in form of a variety of buttery, sweet biscuit crumbs. Nests were enclosed with thin white plastic planks, a smooth material, too slippery for the tarsi of the ants and hence preventing them to forage elsewhere. A square plastic bowl was sunk into the ground and served as feeder. The walls of the feeder were covered with fluon to prevent ants from climbing out. During training ants dropped into the feeder and could return to the nest via a small wooden ramp that led the ants out of the feeder onto the foraging route. Training continued until the ants familiarized with the foraging route and scuttled fast and straight forward between the nest and the feeder at least five times. For tests, ants were either caught at the feeder or close to the nest entrance. Ants caught at the feeder have both the familiar visual scenery and the homing vector of their path integrator as scaffold for homing: hence full-vector ants (FV). Ants caught close to the nest ran off their homing vector and can solely rely on the familiar visual scenery during homing: hence zero-vector ants (ZV). All tested ants were subjected to an eye-cap procedure, which was non-invasive and reversible. For that foragers were manually caught and the first two pair of legs including one of the antenna were carefully fixed between two fingertips. Thus, the head of the ant was immobile and one of the compound eyes could be covered with a drop of opaque enamel paint (Tamiya). The tip of a thin pin acted as brush and painted ants were subsequently checked for an even and complete cover of the targeted eye with the help of a 10× magnifying glass. Afterward ants were transferred into a small plastic vial and tested as soon as the foragers held on to a crumb. Paths of tested ants were recorded with pen and paper and the help of a square grid (0.5×0.5 m) made of string and tent pegs mounted on the ground. Paths were digitized with GraphClick (Arizona Software) and statistical analyses were calculated with Matlab (Mathworks, Matick, MA, U.S.A.).

Paths in the unfamiliar environment (Sham, Just capped and Compensated ants) were recorded with a Panasonic Lumix camera (DMC FZ200) fixed on a tripod, digitized via a novel video tracker system of the Risse group at the University of Münster (52, 53) and analyzed with R studio. LEC and REC ants were pooled by mirroring paths of REC ants to LEC. Differences in their tendency to turn left or right were determined by comparing the angular velocity via a calculation of Dtheta and forward velocity. The X and Y values of the paths were smoothed with a Savitzky-Golay filter of order 3 and filter length of 41 frames, followed by a double smoothing of the Dtheta signal by moving the average of window length by three. Finally, all pauses and events longer than one second were removed. A one-sided Wilcoxon test was used to calculate the significance of each pooled group against random choice.

## ACKNOWLEDGEMENTS

We are grateful for logistic and administrative support of Xim Cerda and his team at the Spanish National Research Council (CSIC Seville) and appreciate the help during field work preparation and data collection of Cornelia Buehlmann, Florent Le Moël and Christelle Gassama. This study was funded by the European Research Council 759817-EMERG-ANT ERC-2017-STG.

