## Supplemental Information for "Compensation to visual impairments and behavioral plasticity in navigating ants"

**This PDF file includes:**

Figures S1 to S3
Table S1

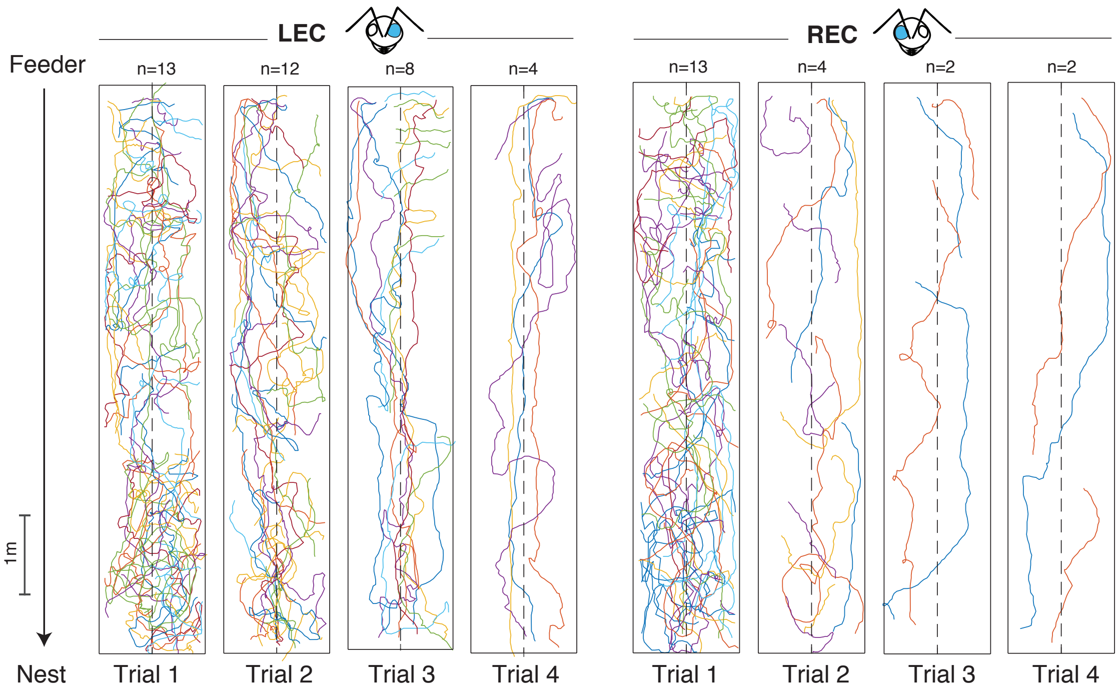


**Fig. S1.** Initial side biases of homing paths in eye capped ants. Black arrows: travel direction of eye capped ants from feeder to nest along a straight foraging route; dashed line: middle of the foraging route; colored lines: single ant paths, gaps between paths occurred when ants ran off the board and were re-released in the middle of the homing route; ant head sketch: eye condition; LEC: left eye covered; REC: right eye covered (this applies to all other supplementary Figures).

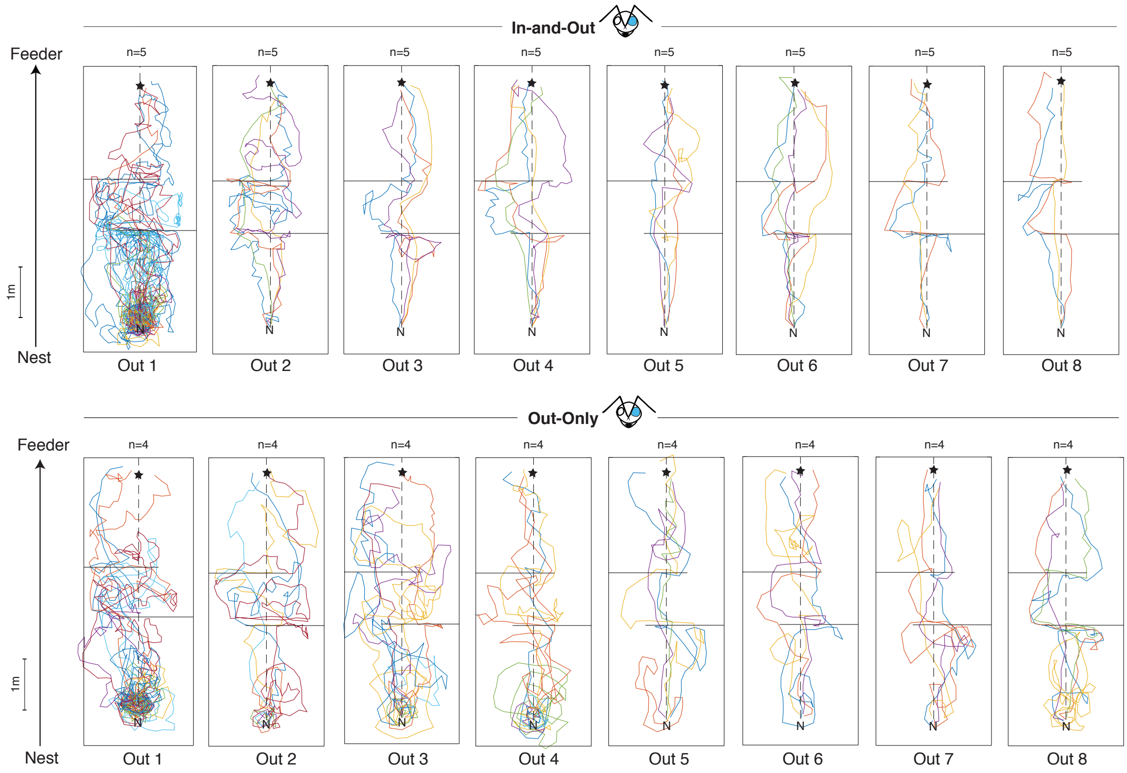


**Fig. S2.** Outbound ontogeny of eye capped ants across trial 1 to 8. Horizontal lines: one-way baffles on the foraging route that could be traversed during outbound trips but had to be negotiated during inbound trips. Left- (LEC) and right (REC) eye capped ants were pooled. Paths of REC ants were mirrored and depicted as LEC ants (painted blue eye in ant head). Only outbound trips are shown. With trial number outbound trips improve and are more directed toward the feeder, especially in In-and-Out ants. Star indicates feeder and N nest position.

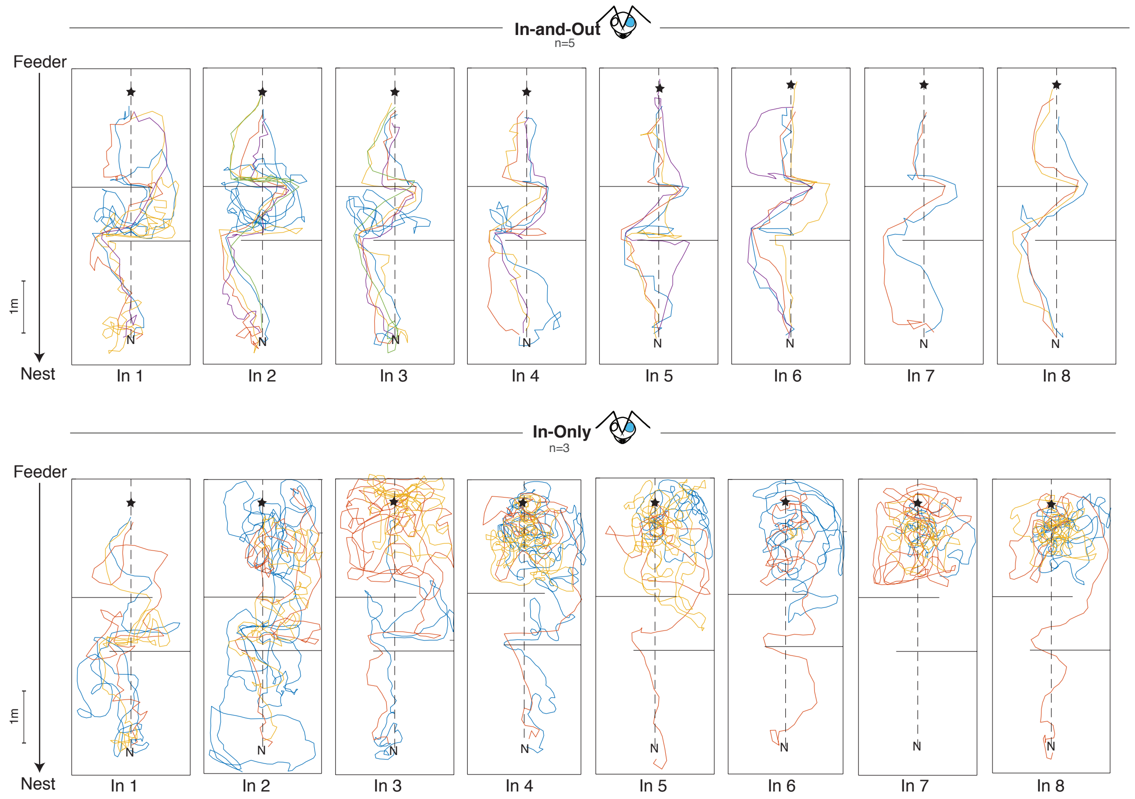


**Fig. S3.** Inbound ontogeny of eye capped ants across trial 1 to 8. Horizontal lines: one-way baffles on the foraging route that could be traversed during outbound trips but had to be negotiated during inbound trips. Left- (LEC) and right (REC) eye capped ants were pooled. Paths of REC ants were mirrored and depicted as LEC ants (painted blue eye in ant head). Only outbound trips are shown. With trial number inbound trips improve and are more directed toward the feeder only In-and-Out ants. In-Only ants struggled to home irrespective of the trial number. Star indicates feeder and N nest position.

**Table S1.** Statistical results of the initial (first 0.2 m) side biases in homing path of eye capped ants (see Fig. 1B) with Rayleigh-test confirming a non-random directional distribution and S-test rejecting nest direction. Significant results are marked in red.

| FV | 1 Trial | 2 Trial | 3 Trial | 4 Trial | Sham |
|---|---|---|---|---|---|
| **Rayleigh-test** | $P = 0.001$ | $P = 0.107$ | $P = 0.026$ | $P = 0.218$ | $P < 0.001$ |
| | $z = 5.601$ | $z = 2.220$ | $z = 3.352$ | $z = 1.584$ | $z = 13.501$ |
| **S-test** | $P = 0.001$ | $P = 0.125$ | $P = 0.023$ | $P = 0.133$ | $P = 0.066$ |
| | $t = -5.218$ | $t = -1.744$ | $t = -3.675$ | $t = -2.460$ | $t = -1.970$ |
| **ZV** | **1 Trial** | **2 Trial** | **3 Trial** | **4 Trial** | **Sham** |
| **Rayleigh-test** | $P = 0.027$ | $P = 0.007$ | $P = 0.031$ | $P = 0.306$ | $P < 0.001$ |
| | $z = 3.333$ | $z = 4.552$ | $z = 3.219$ | $z = 1.264$ | $z = 8.628$ |
| **S-test** | $P = 0.036$ | $P = 0.025$ | $P = 0.014$ | $P = 0.141$ | $P = 0.260$ |
| | $t = -3.102$ | $t = -2.834$ | $t = -4.133$ | $t = -2.374$ | $t = -1.211$ |
